# C-terminal HSP90 Inhibitors Block the HSP90:HIF-1α Interaction and Inhibit the Cellular Hypoxic Response

**DOI:** 10.1101/521989

**Authors:** Nalin Kataria, Bernadette Kerr, Samantha S. Zaiter, Shelli McAlpine, Kristina M Cook

**Affiliations:** University of Sydney, Faculty of Medicine and Health, Charles Perkins Centre, Sydney, Australia; School of Chemistry, University of New South Wales, Sydney, Australia

## Abstract

Hypoxia Inducible Factor (HIF) is a transcription factor activated by low oxygen, which is common in solid tumours. HIF controls the expression of genes involved in angiogenesis, chemotherapy resistance and metastasis. The chaperone HSP90 (Heat Shock Protein 90) stabilizes the subunit HIF-1α and prevents degradation. Previously identified HSP90 inhibitors bind to the N-terminal pocket of HSP90 which blocks binding to HIF-1α, and produces HIF-1α degradation. N-terminal inhibitors have failed in the clinic as single therapy treatments due in part because they induce a heat shock response, which increases chemotherapy resistance. SM molecules are HSP90 inhibitors that bind to the C-terminus and do not activate the heat shock response. The effects of C-terminal HSP90 inhibitors on HIF-1α are unreported. Herein we show that SM compounds block binding between HSP90 and HIF-1α, leading to HIF-1α degradation through the proteasome using the PHD/pVHL pathway in hypoxic conditions. The SM compounds decrease HIF-1α target gene expression at the mRNA and protein level under hypoxia in colorectal cancer cells, leading to cell death, without inducing a heat shock response. Our results suggest that targeting the C-terminus of HSP90 blocks the hypoxic response and may be an effective anti-cancer strategy.

---

Hypoxia is common in tumors and activates a transcription factor known as Hypoxia Inducible Factor-1 (HIF-1). HIF-1 controls the expression of genes involved in glycolysis, angiogenesis and cell survival, and is associated with a poor cancer prognosis^1^. Given that HIF-1 is upregulated in majority of solid tumors, there are ongoing drug development efforts to inhibit its activity. Previous work has focused on inhibiting the HIF-1α subunit from binding to partners such as p300^2–5^ or HIF-1β^6^, or binding to the DNA itself^7^.

HIF-1 is a heterodimer composed of α and β subunits. Activity of the HIF-1 transcriptional complex is regulated post-translationally through degradation of HIF-1α in an oxygen-dependent process^1,8^. HIF-1α stability and degradation is further regulated by the chaperone Heat Shock Protein 90 (HSP90)^9,10^. N-terminal HSP90 inhibitors such as geldanamycin and related analogues bind in the ATP binding pocket on the amino terminus (N-terminus) of HSP90^11^, disassociating it from HIF-1α and inducing pVHL-independent degradation of HIF-1α^9,10^. While N-terminal HSP90 inhibitors effectively block the due to their ability to induce a survival mechanism in cancer cells known as a heat shock response (HSR)^12^. The compounds also have poor selectivity for HSP90^13,14^.

C-terminus inhibitors of HSP90 (SM molecules)^11^ act in a selective manner, and in contrast to the N-terminus inhibitors, they do not induce a heat shock response^13,15^. Although the SM molecules block all co-chaperones that bind to the C-terminus of HSP90 and inhibit HSP90 function^16,17^, there is no data discussing whether C-terminal inhibitors will impact the binding event between HSP90 and HIF-1α or the subsequent hypoxic response. This is the first study examining the effect of C-terminal HSP90 inhibitors on the activity of HIF-1α in hypoxia and downstream gene expression.

Herein we report that the SM compounds prevent accumulation of HIF-1α in HCT116 colorectal cells in hypoxia (0.5% O2) (Fig 1). HIF-1α substantially increases in HCT116 cells under hypoxia. When hypoxic HCT116 cells are incubated in the presence of SM122, HIF-1α protein levels decreased to normoxia levels. SM253 and SM258 also decreased HIF-1α in hypoxia (Fig 1a and 1b). The HSP90 inhibitors were also compared to chetomin, a potent HIF-1 inhibitor that blocks the interaction between HIF-1α and p300 by a zinc ejection mechanism^2^. SM122 reduces HIF-1α levels similar to that seen with chetomin (Fig. 1a and b).

**Fig. 1.**
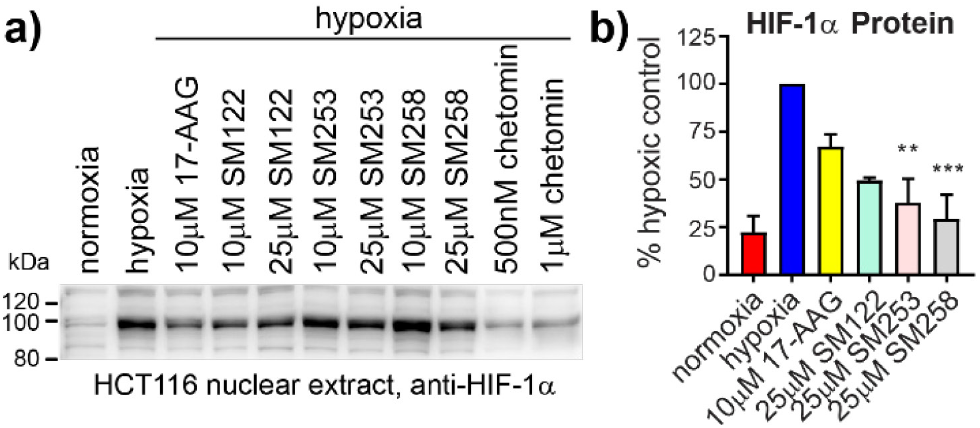
SM compounds inhibit HIF-1α stabilization under hypoxia. a) Nuclear HIF-1α levels in HCT116 cells after 6 h of hypoxia and SM treatment. b) Densitometry of HIF-1α western blots. mean± SEM of 4-5 experiments (except 25 μM SM122 which is n = 2). **P < 0.01, ***P < 0.001, as compared to hypoxic control.

Cells treated with 17-AAG (10 μM, 100 fold over the GI50), an N-terminal HSP90 inhibitor^16,17^ show similar degradation of HIF-1α when treated with SM122 (10 μM), or SM252 (25μM) or SM258 (25μM). The C-terminal inhibitors, SM122, SM253, and SM258, all have a GI_50_ = ~5 μM^16,17^, yet they are highly effective at inducing the degradation of HIF-1α when cells are treated with 2-5 fold over their GI_50_ (Fig. 1a and b), compared to 17-AAG, which is treated at 100 fold over its GI_50_. The considerable amount of HIF-1α degradation in the presence of SM compounds indicates that targeting the C-terminus of HSP90 is a more effective strategy for decreasing HIF-1α than targeting the ATP binding pocket in the N-terminus. HCT116 cells treated with SM compounds showed varying degrees of inhibition of HIF-1 target gene expression in hypoxia based on both mRNA (Fig. 2a) and protein levels (Fig 2b). The expression of carbonic anhydrase IX (CA9) is regulated by HIF-1 and increases under hypoxia (Fig. 2a). Both the N-terminal (17-AAG) and C-terminal compounds (SM) significantly decreased the expression of CA9 mRNA (Fig. 2a), indicating that HIF-1 target gene expression is inhibited by both series. We also examined the mRNA and protein expression of multiple glycolytic genes under HIF-1 control upon exposure to hypoxia and HSP90 inhibitors. At the mRNA level, 25 μM SM258 and 25 μM and 10μM 17-AAG significantly inhibited glucose transporter SLC2A1 (Glut1) mRNA expression (Fig. 2a).

**Fig. 2.**
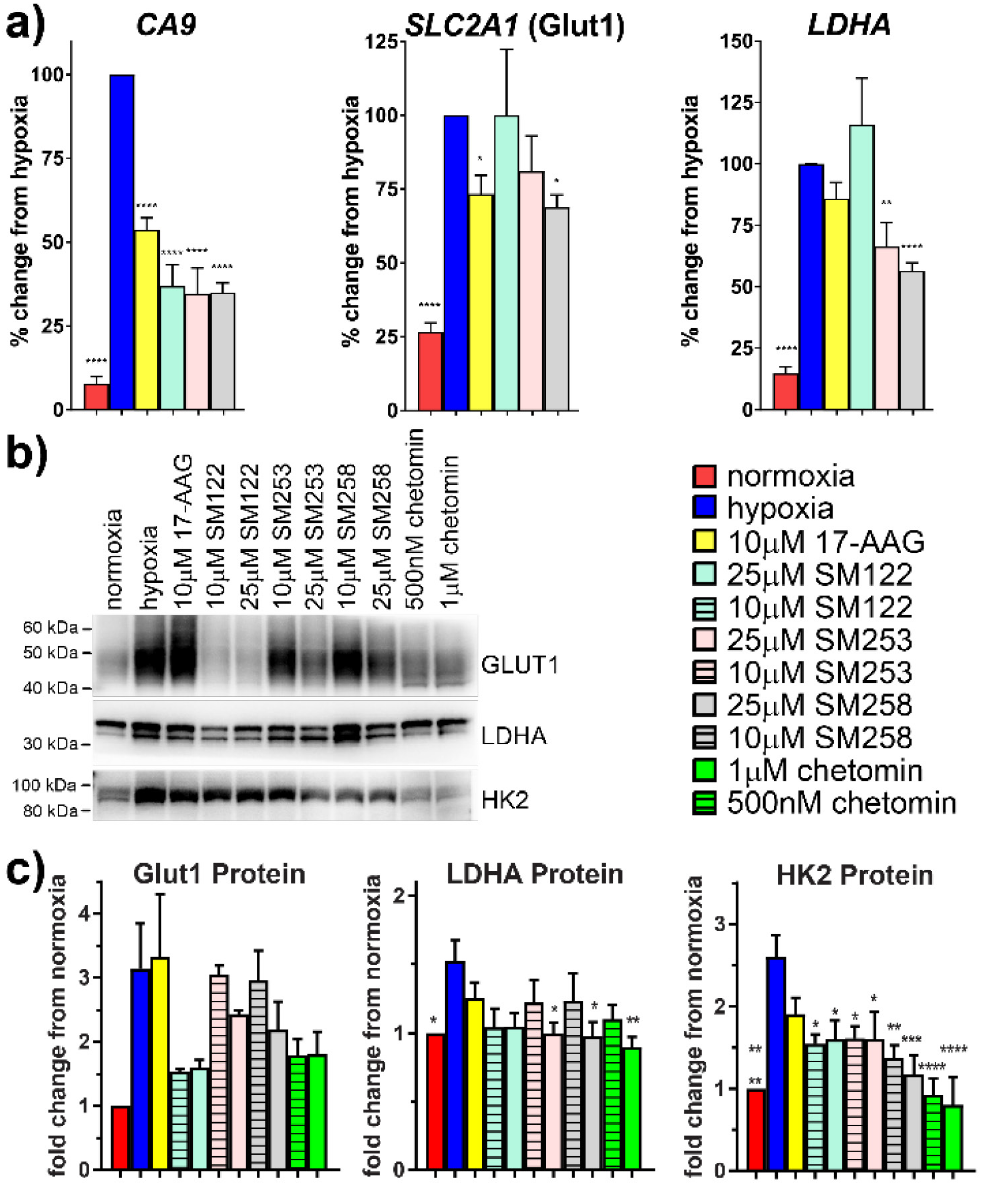
HIF-1 target gene expression is decreased by SM compounds in hypoxia. a) HIF-1 target mRNA expression. HCT116 cells were incubated in the presence or absence of compounds in hypoxia for 18 h. Total RNA was isolated, converted to cDNA and used for qPCR. Results expressed as % of hypoxic control. mean ± SEM (n ≥ 3, except 25 μM SM122 which is n = 2). b) HIF-1 target protein expression. HCT116 cells were incubated in the presence or absence of compounds in hypoxia for 18 h. c) Densitometry of western blots. Results expressed as fold increase over normoxia. (mean ± SEM, n = 5, except Glut1 which is n = 2). *P < 0.05, **P < 0.01, ***P < 0.001, ****P < 0.0001 as compared to hypoxic control.

Comparing the compounds’ impact on Glut1 protein expression, SM122 (25 and 10 μM), produced a significant reduction in the amount of Glut1, similar to levels seen in normoxia (Fig. 2b and c). In contrast, 17-AAG, slightly increased Glut-1 protein levels in hypoxia (10 μM) (Fig 2b and c). Indeed, SM122 was more effective at reducing Glut1 than chetomin (Fig. 2b and c). SM253 and SM258 both reduce Glut1 but cells must be treated at higher concentrations than SM122. These data indicate that SM molecules functionally reduce the glucose transporter and are more potent than the N-terminal inhibitor 17-AAG.

SM253 and SM258 reduced LDHA (lactate dehydrogenase A) mRNA expression under hypoxia (Fig 2a). SM122 had little effect on LDHA mRNA. SM253 and SM258 both decreased LDHA protein to similar levels as chetomin (~33%) (Fig 2b and c). Protein levels of hexokinase 2 (HK2), another enzyme involved in glycolysis, were significantly reduced by SM122 and SM253 (~40%) (Fig. 2b and c). SM258 decreased HK2 by ~60%. Thus, SM122 was highly effective at reducing HIF-1 target protein expression for Glut1, LDHA and HK2 compared to N-terminal HSP90 inhibitor 17-AAG, whilst SM253 and SM258 were slightly more effective than 17-AAG at reducing Glut1. This indicates that C-terminal SM compounds, and particularly SM122, are effective at reducing glycolytic enzymes that are commonly upregulated in cancers.

To test the cytotoxic effect of SM compounds, we measured HCT116 cell viability at 18 and 72 h and compared to 17-AAG. The majority of cells were still viable at concentrations up to 25 μM at 18 h (see supplementary), where HIF-1 inhibition was documented (Fig. 2). However, after 72 h, cell viability significantly decreased under normoxia and hypoxia for all drugs indicating HSP90 is essential for cell survival.

A pitfall of N-terminal HSP90 inhibitors is that they induce a heat shock response (HSR). The HSR activates heat shock factor-1, a transcription factor that controls expression of HSP90, 70, 40 and 27^18^. The HSR therefore increases levels of HSP90, the protein targeted by N-terminal drugs, and activates mechanisms that block apoptosis and increase chemo-resistance^18^. SM HSP90 inhibitors do not induce the HSR in normoxia thus greatly increasing their cellular efficacy^13^. The HSR under hypoxic conditions is unknown, as is the impact of SM compounds on HSP protein levels.

To test if a heat shock response is activated by N- and C-terminal HSP90 inhibitors in hypoxia, we measured HSP related proteins following an 18 h drug treatment. Treatments using 10 μM 17-AAG produced high levels of HSP70, and a slight increase in HSP90, while the SM compounds did not (Fig. 3a and b). We also compared the heat shock response from the HSP90 inhibitors to chetomin (Fig. 3a and b). At 1 μM chetomin, HSP70 significantly increases. This is not entirely unexpected given that the heat shock response can be induced by cell stress and chetomin is associated with associated with significant cytotoxicity^2,5^.

**Fig. 3.**
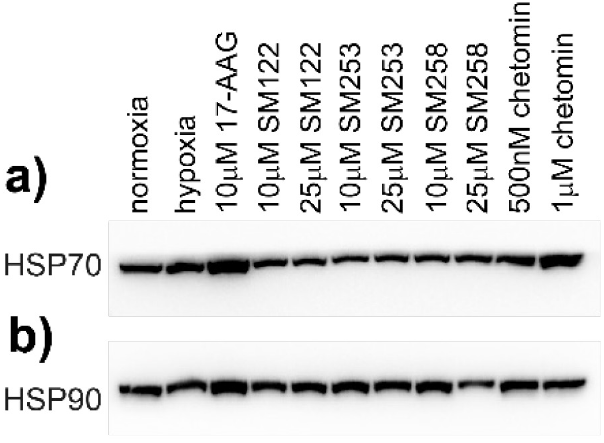
3 SM compounds do not induce a heat shock response like the N-terminal inhibitor 17-AAG. HCT116 cells were incubated for 18 h in the presence or absence of compounds in hypoxia and compared to normoxia (n=3-4).

Under normoxic conditions, HIF-1α is degraded by the proteasome. Prolyl hydroxylase enzymes (PHDs) hydroxylate HIF-1α using oxygen as a substrate, which enables binding of pVHL and the E3 ubiquitin ligase complex, leading to proteasomal degradation of HIF-1α. To test if HIF-1α was being degraded by the proteasome in hypoxia, we used a proteasome inhibitor, MG262, in combination with the SM compounds in hypoxia. Treating HCT116 cells with MG262 in hypoxia leads to an increase in HIF-1α accumulation in the cytoplasm and nucleus (Fig. 4a, b and c). This increase is clearly visible in cytoplasmic HIF-1α as proteasomal degradation occurs in the cytoplasm and is inhibited by MG262. Treatment with SM258 alone caused significant degradation to HIF-1α in the cytosol. SM122 and SM253 were as effective as 17-AAG in degrading cytosolic HIF-1α (Fig. 1 and 4a and b). Co-treatment of SM compounds with MG262 increased nuclear and cytosolic HIF-1α accumulation in hypoxia, suggesting that the proteasome is involved in degrading HIF-1α when the C-terminus of HSP90 is inhibited (Fig. 4a, b and c).

**Fig 4.**
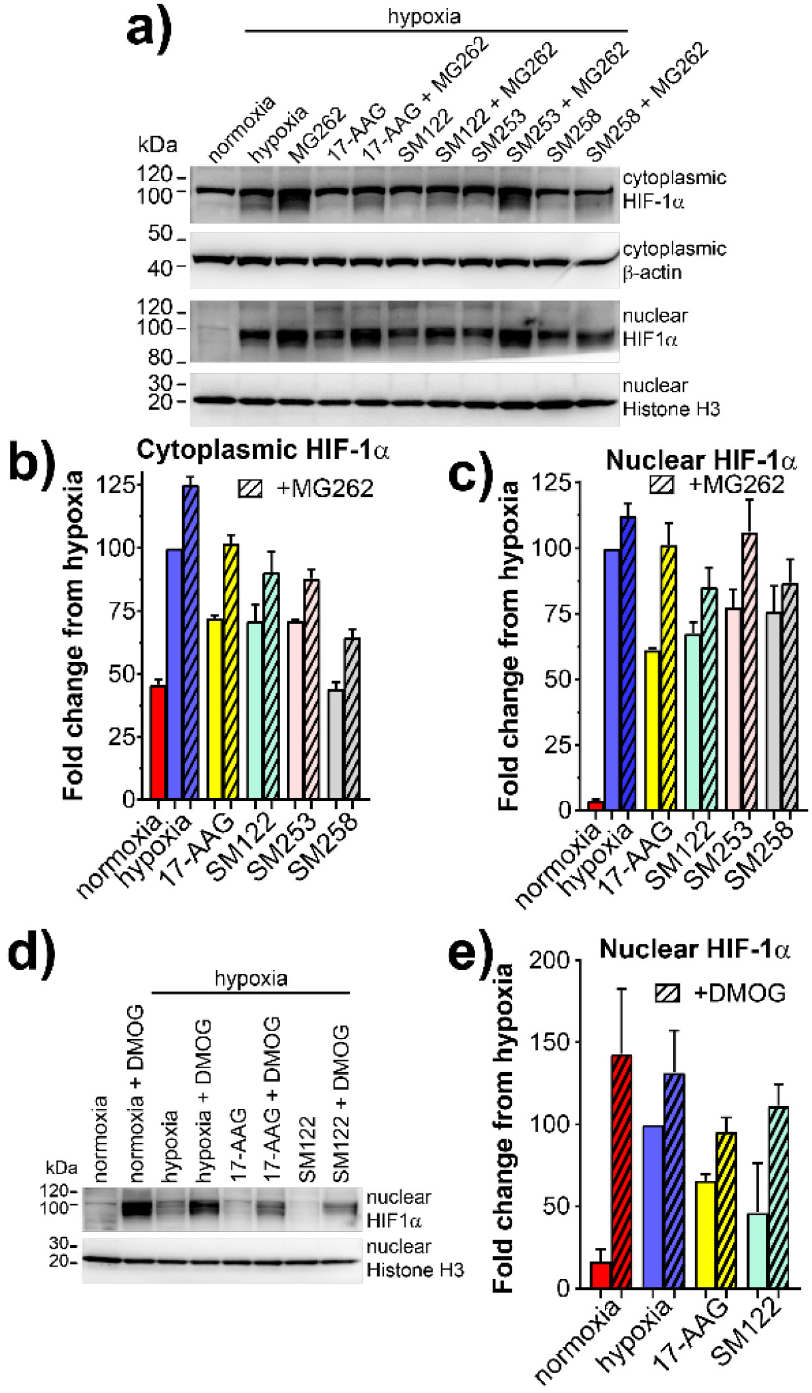
SM compounds degrade HIF-1α via the PHDs and proteasome in hypoxia. a) HCT116 cells were treated with HSP90 inhibitors (25 μM) and/or the proteasome inhibitor MG262 (1 μM) for 6 h in hypoxia. b) Densitometry of cytoplasmic HIF-1α (b) and nuclear HIF-1α (c) (n = 2). d) HCT116 cells were treated with 25 μM SM122 and/or 1 mM DMOG, which inhibits the PHD prolyl hydroxylases, for 6 h in hypoxia. e) Densitometry of nuclear HIF-1α (mean±SEM, n = 2).

To determine if the PHDs were involved in degradation of HIF-1α by the SM compounds, we used a PHD inhibitor, dimethyloxalylglycine (DMOG) in hypoxia. Surprisingly, treating HCT116 cells with DMOG in hypoxia leads to a slight increase in HIF-1α accumulation in the nucleus (Fig. 4d). This indicates that PHD-mediated hydroxylation occurs even at very low levels of oxygen, such as the 0.5% used for our experiments. Treating cells with the HSP90 inhibitors, 25 μM 17-AAG or 25 μM SM122, decreases the amount of HIF-1α under hypoxia, while combining either drug with DMOG increases the amount of HIF-1α in the nucleus under hypoxia, particularly with SM122 treatment (Fig. 4d and e). This means that PHD activity is required for HIF-1α to be degraded by the proteasome in the presence of both our C-terminal HSP90 inhibitor, SM122 and the N-terminal inhibitor, 17-AAG.

N-terminal HSP90 inhibitors, such as geldanamycin, possess anti-angiogenic activity by inhibiting HIF-1α which controls expression of pro-angiogenesis genes^1^. HSP90 also interacts with other pro-angiogenic proteins, preventing their stabilization and contributing towards inhibition of angiogenesis^19^. To test the anti-angiogenic effects of the C-terminal SM compounds, an endothelial cell tube formation assay was used. After coating plates with a basement membrane extract, HUVECs were added with HSP90 inhibitors. In the presence of vehicle only (1% DMSO), an extended network of tubules formed (Fig. 5a). SM122 (25 μM) was highly effective at inhibiting tubule formation and decreasing both the total area and the number of meshes. SM253 (25 μM) was also effective, followed by 10 μM 17-AAG. SM258 (25 μM) did not appear to impact on tubule formation. Our results indicate that C-terminal HSP90 compounds, SM122 and SM253, possess anti-angiogenic activity in endothelial assays. Both HIF and HSP90 play a role in angiogenic processes^1^ and inhibiting this interaction decreases endothelial tubule formation.

**Fig. 5.**
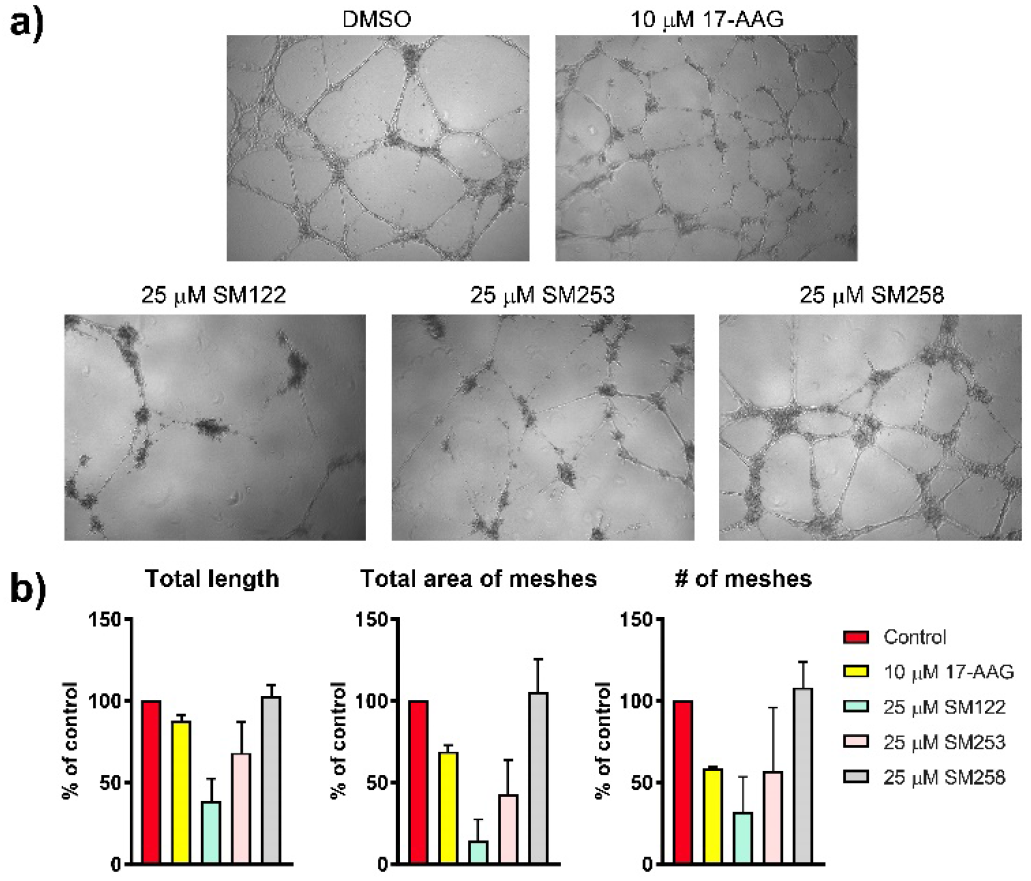
SM compounds decrease endothelial tube formation. a) HUVEC cells were treated with inhibitors and plated on EHS matrix. Images taken after 18 h. Representative images shown. b) Tubule formation was quantified using ImageJ with the add-on Angiogenesis Analyzer (NIH) (n = 2).

This is the first comprehensive study of how a new series of C-terminal HSP90 inhibitors, SM122, SM253 and SM258, impact the HIF hypoxic response. These molecules lead to HIF-1α degradation and decrease downstream HIF-1 target gene expression. Specifically, Glut1 (SLC2A1) imports glucose and increased expression is associated with decreased cancer patient survival^20^. Glut1 itself has been investigated as a cancer drug target in order to block glucose uptake and limit glycolysis. Treating colorectal cancer cells with SM molecules, specifically SM122, reduced Glut1 expression on the mRNA and protein level, thereby impacting tumour cell behavior (Fig. 2). The SM compounds were generally more effective HIF-1α inhibitors compared to the N-terminal HSP90 inhibitor, 17-AAG, which induces a heat shock response (HSR) in both normoxia and hypoxia in HCT116 cells. In contrast, the C-terminal SM compounds do not induce a heat shock response (Fig. 3). This is significant given that the heat shock response from N-terminal inhibitors directly counteracts the efficacy of the HSP90 inhibitors and is thought to be a primary factor in the failure of HSP90 inhibitors in the clinic^12,21^. Decreasing the heat shock response using HSP siRNAs significantly increases the potency of the N-terminal HSP90 inhibitors^22,23^. The SM series overcomes this hurdle by not inducing the heat shock response.

The SM compounds also degrade HIF-1α in hypoxia through the proteasome (Fig. 6). The PHDs and canonical oxygen degradation pathway is still relevant for HIF-1α degradation by the SM compounds even in hypoxic conditions as low as 0.5% oxygen (4 mmHg). While somewhat surprising that the PHDs remain active under low oxygen conditions, the genes encoding the three PHD enzymes (EGLN1, EGLN2 and EGLN3) are expressed by HIF-1 itself^24^, meaning that the amount of available PHD enzymes significantly increases in hypoxia and act as a feedback loop to limit an excessive hypoxic response. PHD activity has been observed at oxygen levels as low as 0.2%^24^, which is significantly lower than the conditions we used. The reduction of endothelial tube formation (Fig. 5) supports the effective anti-angiogenic mechanism by which the C-terminal inhibitor, SM122, functions.

**Fig 6.**
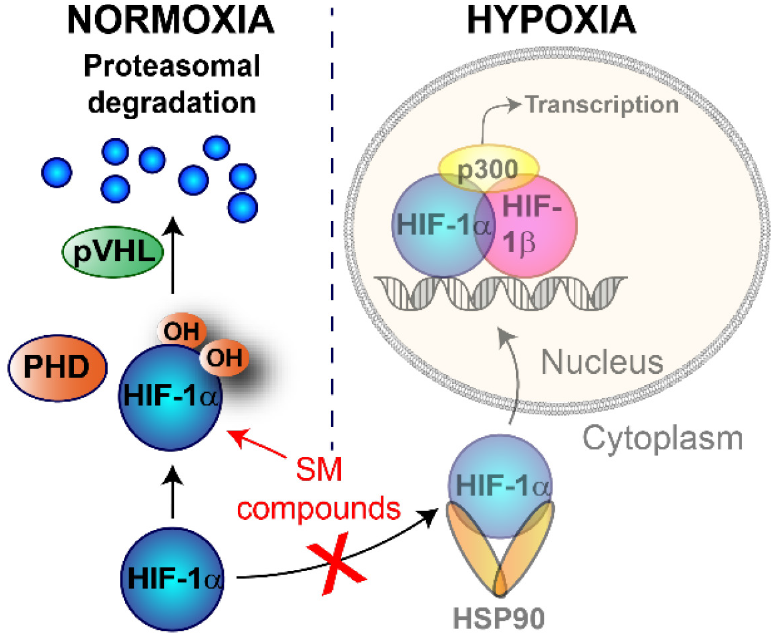
Mechanism of HIF-1α degradation by SM compounds. a) Regulation of HIF-1α by oxygen. Under normal oxygen conditions, HIF-1α is hydroxylated by the prolyl hydroxylases (PHDs), which uses oxygen as a substrate. Hydroxylation by the PHDs enables pVHL and the E3 ubiquitin ligase complex to bind HIF-1α, signaling for proteasomal degradation. Under hypoxic conditions, hydroxylation decreases enabling HSP90 to stabilize HIF-1α and, allowing formation of the active HIF-1 complex. Under hypoxic conditions in the presence of SM compounds, HSP90 is inhibited, destabilizing HIF-1α. This allows the PHDs to hydroxylate HIF-1α, even under low oxygen conditions, targeting it for proteasomal degradation

Thus, using SM compounds that target the HSP90 C-terminus is an effective strategy for blocking HIF-1 activity and reducing both tumor growth and angiogenesis. SM compounds are a promising anti-cancer strategy for targeting HSP90 and impacting HIF-1α function.

## Methods

### Cell culture

HCT116 cells were maintained in McCoy’s 5A medium (Gibco), supplemented with 10% fetal bovine serum (FBS), 50 μg/ml Streptomycin, 50 IU/ml Penicillin and 1x GlutaMAX. For non-hypoxic experiments, cells were incubated in humidified air supplemented with 5% CO2 at 37°C (18.6% O2 v/v). Cell lines were a gift from Professor Philip Hogg and from ATCC, Bethesda, MD. Cell lines are tested four times per year for mycoplasma contamination.

### Cell viability assays

For 72 h viability assays, HCT116 cells were seeded into 96-well plates in 100 μl of medium at a concentration of 2 × 103 cells/well (for 18 h viability assays 5.1 × 104 cells/well were used). After overnight incubation at 37 °C, medium was removed and replaced with 100 μl of fresh medium (with 0.5% fetal bovine serum) containing drug or vehicle control (1% DMSO). Plates were placed in either a normoxic (18.6% oxygen) or hypoxic (0.5% oxygen) incubator for 18 or 72 h. Cell viability was measured by adding 20 μl CellTiter-Blue cell viability reagent (Promega) to each well, after which the cells were returned to the 37 °C incubator until sufficient color change. Fluorescence intensity was measured using 570 nm excitation and 600 nm emission on a Tecan Infinite M1000 Pro.

### Western blots

HCT116 cells were seeded into 12 well plates at 6 × 105 cells/well. The following day, cells were treated with medium containing SM compounds or vehicle control (DMSO). Plates were placed in incubators under normoxic or hypoxic conditions for 6 – 18 h. Cells were lysed using RIPA buffer containing protease and phosphatase inhibitors (Roche). Lysates were sonicated, mixed with 4x LDS sample buffer (ThermoFisher Scientific) containing DTT and heated to 70 °C for 10 min.

SDS-solubilized protein samples were resolved using the Novex NuPage SDS-PAGE gel system (ThermoFisher Scientific; 4-12% Bis-Tris gels), and transferred via semi-dry electrophoresis to 0.45 μm PVDF membranes (Millipore). Membranes were blocked for 1 h in 5% non-fat dry milk in TBST and then incubated overnight at 4°C in one of the following antibodies: anti-GLUT1 (ab115370, 1:1000, Abcam), anti-HIF1α (NB100-479, 1:500, Novus), anti-Histone H3 (ab1791, 1:2000, Abcam), anti-beta Actin (1:2000, ab8226, Abcam), anti-HK2 (TA325030, 1:500, Origene), anti-LDHA (3582T, 1:1000, Cell Signaling Technologies), anti-HSP90 (NB120-1429, 1:1000, Novus) and anti-HSP70 (4873, 1:1000, Cell Signaling Technologies). All blots used Histone H3 as a housekeeping protein to ensure equal loading. Following washes, blot was incubated for 1 hour room temperature in a 1:1000 dilution of corresponding secondary antibodies (Dako). Bound antibodies were visualized using the BioRad Chemidoc Touch System. Densitometry was calculated using BioRad ImageLab software. For Fig 3, blots were cut horizontally between 30 kDa - 40 kDa markers, with the top portion probed for either HSP90 or HSP70 and the bottom probed for Histone H3 to ensure equal loading (not shown). The HSP90 antibody does not differentiate between the HSP90β1 (HSP90B1) and HSP90α (HSP90AA1) isoforms, so total HSP90 protein is reflected in the blot. The HSP70 antibody reacts with the inducible protein HSP70-1A (HSPA1A).

For Fig 4, HIF-1α western blots, the blot was horizontally cut to probe proteins between ~80 kDa – 220 kDa, as specified by the antibody manufacturer. The lower portion of the blot was used to probe for the housekeeping proteins, Histone H3 or β-actin. Cytoplasmic HIF-1α and β-actin in (a) were cut from the same gel/blot. Nuclear HIF-1α and Histone-H3 in (a) were cut from the same gel/blot. The blots for nuclear HIF-1α and Histone-H3 in (d) were cut from the same gel/blot.

### Treatment with proteasome or PHD inhibitors

HCT116 cells were seeded into 12-well plates at 3 × 105 in 1 ml of McCoy’s media with 10% FBS and glutamax. After overnight incubation at 37 °C, medium was removed and replaced with 1 ml of fresh medium (with 0.5% fetal bovine serum) containing SM compounds and drug (1 mM DMOG for PHD inhibition or 1 mM MG-262 for proteasome inhibition) or vehicle control (1-2% DMSO). Plates were placed in either a normoxic (18.6% oxygen) or hypoxic (0.5% oxygen) incubators for 6 h before harvesting as described for mRNA or protein analysis. DMOG (dimethyloxalylglycine) was purchased from Sigma Aldrich and MG-262 was purchased from Calbiochem.

### Nuclear and cytoplasmic protein separation

Nuclear and cytoplasmic fractions were separated using the NE-PER Nuclear and Cytoplasmic Extraction Reagents kit (ThermoFisher Scientific) according to the manufacturer’s instructions.

### Gene expression

To determine if the SM compounds decrease HIF-1 target gene expression, we exposed HCT116 cells to SM compounds in hypoxia (0.5% O2) for 18 h, which is the approximate time scale for maximum HIF-1 target mRNA expression^25^. Total RNA was isolated using the RNeasy Mini Kit (Qiagen). RNA quality was assessed using an RNA bleach gel according to^26^. The quantity of isolated RNA was measured using a Nanodrop (260nm). Total RNA (2 μg) was used as a template for cDNA synthesis using the High-Capacity cDNA Reverse Transcription Kit (Applied Biosystems). For qRT-PCR, pre-designed TaqMan Gene Expression Assays were used with TaqMan Gene Expression Master Mix (Life Technologies, Thermo-Fisher). 20 ng of cDNA was used per well/reaction. 18s RNA was used to normalize samples. qRT-PCR protocol followed according to manufacturer on a QuantStudio 7 Flex machine. Fold-change in mRNA levels was calculated using the ΔΔCt method. TaqMan Primers (Assay ID) used for qPCR (ThermoFisher Scientific) were as follows: CA9 (carbonic anhydrase IX) (Hs00154208_m1), SLC2A1 (Glut1) (Hs00892681_m1), LDHA (Hs01378790_g1), PGK1 (Hs00943178_g1), IGFBP3 (Hs00181211_m1), PLOD2 (Hs01118190_m1), HSP90B1 (HSP90B1) (Hs00427665_g1), HSPA1A (HSP70-1A) (Hs003 59163_s 1), HSPA8 (Hsc70) (Hs03044880_gH), HSP90AA1 (HSP90α) (Hs00743767_sH), HSPB1 (HSPβ1 or HSP27) (Hs00356629_g 1), RACK1 (Hs00272002_m1) and 18s (Hs03003631_g1).

### Endothelial cell tube formation assay

HUVEC cells were grown in EGM-2 media (Lonza, Walkersville USA) as before^27^. For tube formation, 96 well plates were treated with 38 μl EHS Matrix Extract (Sigma-Aldrich) for 60 min at 37° C. HUVECs (24,000 cells per well) were plated on top and treated with 25 μM 17-AAG, 10 μM 17-AAG, 25 μm SM122, 25 μM SM253 or 25 μM SM258. DMSO was used as a control. Plates were incubated for 18 h and imaged using phase contrast microscopy. The program ImageJ with the Angiogenesis Analyzer (National Institutes of Health) was used to analyze images of tubule formation. The program was set to calculate total segment length, total area of meshes and the number of nodes. Three iterations were used with the minimum object size set to 50 pixels. The master segment threshold was set to 30 pixels, with the minimum branch size at 25 pixels, artifactual loop size set to 1000 pixels and the isolated element threshold set to 100 pixels.

SM Compounds solid-phase peptide synthesis Stepwise SPPS was performed in a polypropylene solid-phase extraction cartridge fitted with a 20 μm polyethylene frit purchased from Applied Separations (Allentown, PA).

### Coupling reaction and FMOC removal

Prior to each coupling reaction, the resin was swelled in DMF for 0.5 – 1 h, then the DMF was drained. Couplings were performed in DMF at a concentration of 0.3 M. Fmoc-protected amino acid (2 equiv.) and either 1-Hydroxybenzotriazole hydrate (HOBt) or 1-Hydroxy-7-azabenzotriazole HOAt (2 equiv.) were mixed with the resin. N, N′-Diisopropylcarbodiimide (DIC) (4 equiv.) was then added to activate the reaction. Coupling reaction was run for a minimum of 2 hours while shaking (Labquake tube shaker, Thermo Fisher Scientific) at room temperature. A negative ninhydrin test was used to confirm reaction completion. Once completed, the reaction mixture was drained, and the resin was subjected to Fmoc Removal. The Fmoc protecting group was removed using the following washes: DMF (3 x 1 min), 20% piperidine in DMF (1 x 5 min), 20% piperidine in DMF (1 x 10 min), DMF (2 x 1 min), iPrOH (1 x 1 min), DMF (1 x 1 min), iPrOH (1 x 1 min) and DMF (3 x 1 min). The resin was then ready for the next coupling reaction.

### Resin cleavage of linear peptide

Once the desired peptide was generated, the final Fmoc protecting group was removed following Fmoc Removal procedure with the following additional washes: DMF (3 x 1 min), iPrOH (3 x 1 min) and MeOH (3 x 1 min). The resin-bound peptide was then dried in vacuo overnight. The resin was then cleaved from the linear peptide using TFE and CH2Cl2 (1:1 v/v) at a concentration of 10 mL/g resin. The reaction mixture was stirred at room temperature for 24 hours before filtering the resin. The filtrate was concentrated and washed at least 10 times with CH2Cl2 to remove residual entrapped TFE. The product was then dried in vacuo overnight to produce the linear peptide.

### Macrocyclisation

Macrocyclisation of the linear peptide was achieved using a cocktail of 3 coupling reagents: HATU (1 eq.), TBTU (0.8 equiv.) and DMTMM (0.8 equiv.). A special case occurred where macrocyclisation was achieved using only HATU (1 equiv.) and TBTU (1 equiv.). DMTMM was removed as the methylmorpholinium by-product from the cyclisation formed a stable complex with the peptide and this by-product was inseparable by HPLC and was only detected via NMR. The reaction was performed under nitrogen and in dilute conditions using anhydrous solvents at concentration of 0.001 M. The linear peptide and coupling reagents were dissolved separately in CH2Cl2, where 20% of the final volume was used to dissolve the linear peptide and the other 80% dissolved the coupling reagents. DIPEA (4 equiv.) was added to each solution. The linear peptide solution was then added drop-wise to the coupling reagents solution via a syringe pump over approximately 2 hours. The reaction was stirred overnight and monitored via LC/MS. (Note: if the reaction failed to reach completion after stirring overnight, additional HATU (1 equiv.) was added and the reaction was monitored using LC/MS.) Upon completion, the reaction mixture was evaporated and the dry solid was redissolved in CH2Cl2 and extracted 3 times with milli-Q water. The aqueous layers were combined and extracted 3 times with fresh CH2Cl2. All organic layers were combined and dried over Na2SO4¬, filtered and evaporated under reduced pressure before the compound was dried in vacuo overnight.

### Statistics

Data are expressed as mean ± SEM from independent experiments. Statistical analysis was performed using a One-Way ANOVA function with multiple comparisons followup test (comparing the mean of each column with the mean of every other column). *P < 0.05, **P < 0.01, ***P < 0.001, ****P < 0.0001, ns = not significant. Figure specific statistical tests: Fig 1: One-Way ANOVA function with multiple comparisons followup test (compare the mean of each column to the hypoxic mean). Fig 2: One-Way ANOVA function with multiple comparisons follow-up test (compare the mean of each column to the hypoxic mean).

## Notes and references

Supplementary data are in Supporting Information.

## SUPPLEMENTARY FIGURES AND DATA

**Fig S1.**
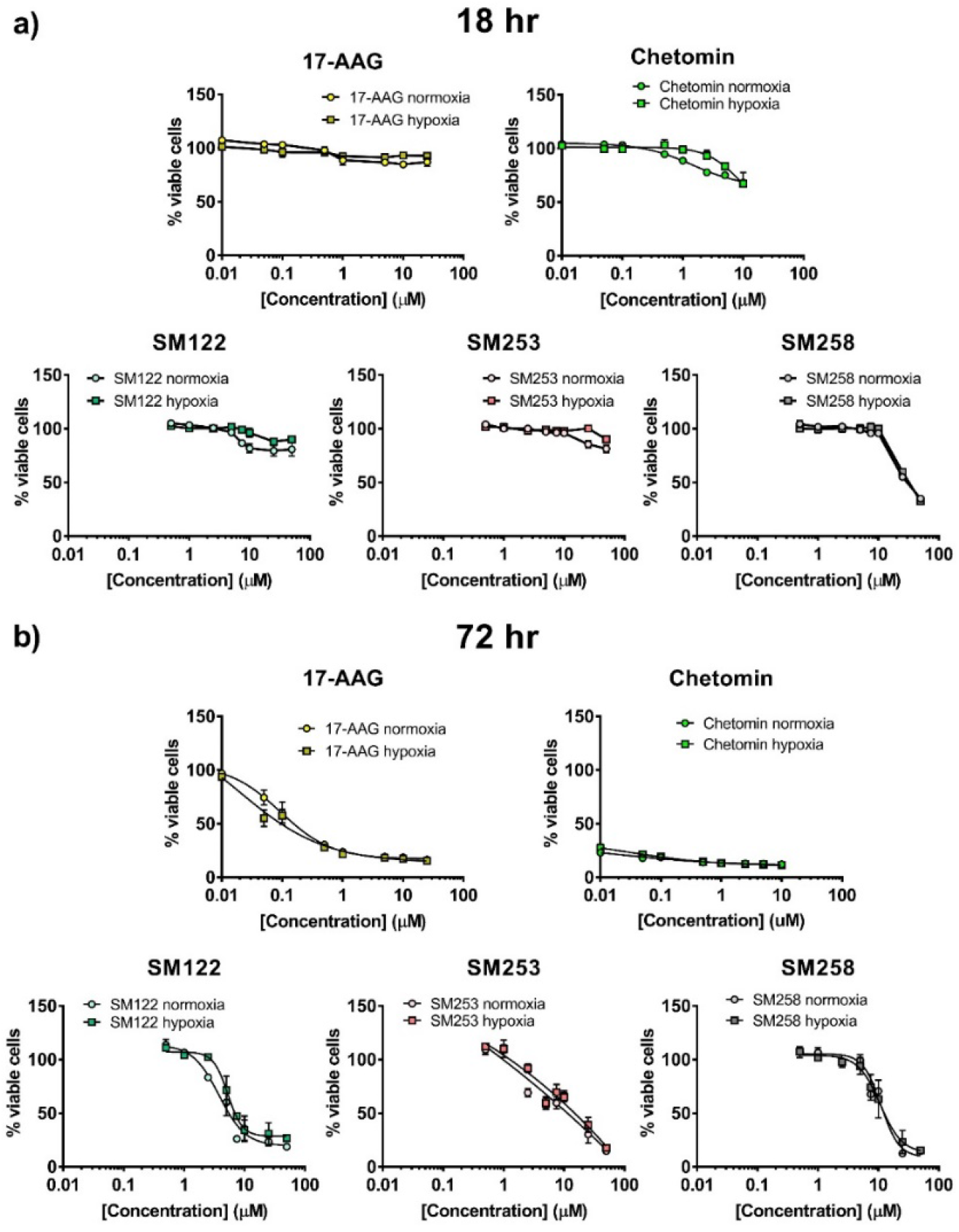
Cytotoxicity of SM compounds, 17-AAG and chetomin in normoxia and hypoxia. The effect of SM compounds on cell viability effects are similar in normoxia and hypoxia, indicating that SM compounds may be an effective strategy for targeting hypoxic tumor cells which are typically more resistant to chemotherapy. HCT116 cells were treated with SM compounds, 17-AAG and chetomin and incubated in normoxic or hypoxic conditions for 18 h **(a)** or 72 h **(b)** before adding Celltiter Blue Cell Viability reagent to assess metabolic capacity. Percent viable cells was normalized to DMSO treated control cells (100%). Results are the mean ± S.E.M of independent experiments run in duplicate (n = 3).

**Fig S1 Results.** The SM compounds initially show little cytotoxicity at 18 h, i.e. HIF-1 inhibitory doses, but produce cell death after 72 h incubations as essential cell survival pathways succumb to complete inhibition over the extended time. The observed cytotoxicity was similar for cells treated under both normoxic and hypoxic conditions. The HSP90 inhibitors were compared to chetomin which is known to have significant cytotoxic and necrotic effects, possibly independent from its effects on HIF-1α. Chetomin was significantly more cytotoxic than the SM compounds and at concentration below HIF-1 inhibitory doses, while the SM compounds show cytotoxicity mainly at the concentrations that inhibit HIF-1 activity. These data indicate that the SM compounds are likely inducing cytotoxicity via a HIF-1 related mechanism, e.g. through inhibition of HIF-1 interacting with HSP90.

